# Spectral brain signatures of aesthetic natural perception in the alpha and beta frequency bands

**DOI:** 10.1101/2022.08.11.503584

**Authors:** Daniel Kaiser

## Abstract

During our everyday lives, visual beauty is often conveyed by sustained and dynamic visual stimulation, such as when we walk through an enchanting forest or watch our pets playing. Here, I devised an MEG experiment that mimics such situations: Participants viewed 8s videos of everyday situations and rated their beauty. Using multivariate analysis, I linked aesthetic ratings to (1) sustained MEG broadband responses and (2) spectral MEG responses in the alpha and beta frequency bands. These effects were not accounted for by a set of high- and low-level visual descriptors of the videos, suggesting that they are genuinely related to aesthetic perception. My findings provide a first characterization of spectral brain signatures linked to aesthetic experiences in the real world.

## Introduction

During our everyday lives, we often encounter beauty in our visual surroundings. Such experiences are not exclusive to stimuli designed to evoke beauty, like artworks or architecture; beauty can also be found in seemingly mundane everyday experiences, like enjoying nature or observing animals. What happens in our brains during such experiences?

Research in neuroaesthetics has made great progress by studying responses to static naturalistic images, such as pictures of faces [1–3] and scenes [4–6]. However, recent investigations suggest that dynamic, compared to static, visual stimuli can evoke stronger feelings of beauty and greater task engagement [7].

So far, only few studies focused on such dynamic aesthetic experiences. A recent fMRI study [8] showed that beautiful dynamic scenes activate the hippocampus more strongly than beautiful static scenes. Another study reported activation changes in the default-mode and fronto-parietal attention networks when participants watched awe-inducing videos [9]. Interestingly, Isik and Vessel [10] showed that the beauty of dynamic visual landscapes is represented in regions distinct from regions activated by static landscapes [6], suggesting that dynamic aesthetic experiences lead to distinctive brain dynamics.

One possibility is that dynamically evolving natural stimuli give rise to unique spectral brain signatures. Our brains exchange information across regional neural assemblies through oscillatory codes [11,12], where brain regions couple or de-couple as a function of external inputs and cognitive states. Studying sustained aesthetic experiences with dynamic stimuli provides a unique opportunity for uncovering how spectral responses relate to aesthetic perception.

Here, I tested whether aesthetic experiences from rich and dynamic natural stimuli are indeed accompanied by characteristic spectral brain dynamics. Using multivariate analyses of MEG data, I found that neural activations in the alpha and beta frequency range predicted participants’ ratings, providing a spectral brain signature for aesthetic perception in dynamic real-world context.

## Materials and Methods

### Participants

Twenty adult volunteers with normal vision participated (mean age 24.2, SD=7.0; 16 female, 4 male). They received monetary reimbursement or course credits. Sample size was constrained by scan time availability. With 20 participants, medium-to-large effects sizes greater than d=0.57 (one-sided t-test) can be detected with 80% statistical power. All participants provided written informed consent. Procedures were approved by the Research Ethics Committee of the York Neuroimaging Centre and adhered to the Declaration of Helsinki.

### Stimuli and Paradigm

Stimuli were 75 8s-videos (1280 × 720, 30Hz) of everyday events, spanning five categories (15 each): transport, animals, forests, water, mountains (Fig. 1a/b). Videos were downloaded and cut from Youtube.com by the author. In each of four runs (~15min each), all videos (17×9.6deg visual angle) were shown in random order. Participants were instructed to maintain fixation and rate how aesthetically pleasing, or beautiful, they found the video using a 4-level circular rating scale (Figure 1a). The rating screen appeared after the video presentation, and participants were allowed to take as much time as they needed to provide their response. The position of response options varied (always increasing clockwise) to disentangle ratings from motor responses. To assess the stability of beauty ratings across the four video repetitions, I performed split-half reliability analyses, where each participant’s ratings across videos from half of the runs were correlated with ratings from the other half of runs; correlations were averaged across all possible splits. Across participants, this revealed a good average reliability of r=0.85 (SE=0.02).

**Figure 1.**
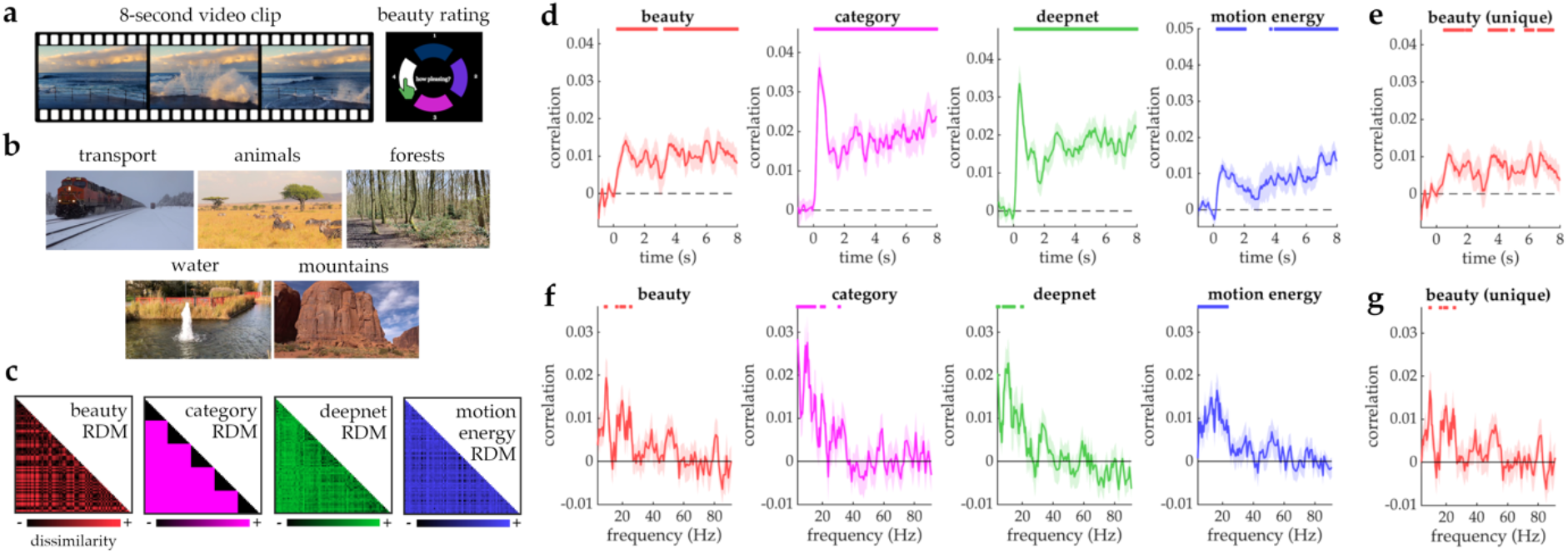
**a)** Illustration of the rating paradigm. **b)** Examples from the 5 video categories. **c)** Neural similarities (across time or frequency) were modelled with similarities in beauty ratings, category, deepnet features, and motion energy. **d)** Correlations between broadband responses and representational models. **e)** Correlation between broadband responses and beauty ratings when the other predictors were partialed out. **f)** Correlations between spectral responses and representational models. **g)** Correlations between spectral responses and beauty ratings when the other predictors were partialed out. Error bars represent SEM. Significance markers denote p_corr_ <0.05.

### MEG recording and processing

MEG was recorded at 1000 Hz using a 248-channel 4D Neuroimaging Magnes3600 system. Preprocessing was performed using the fieldtrip toolbox for Matlab [13]. Data were downsampled to 200Hz, epoched from −1s to 9s relative to onset, and baseline-corrected. Noisy channels were removed and interpolated using neighboring channels. Eye and heart artifacts were removed using ICA.

### Spectral responses

For computing spectral responses, I split every trial into 4 bins of 2 seconds (0-2s, 2-4s, 4-6s, 6-8s), which were treated as individual samples. For each channel and sample, powerspectra were obtained from DPSS multitaper analyses [13] with spectral smoothing of ±3Hz (4-30Hz) or ±6Hz (31-91Hz). Power values were computed in 0.5Hz-steps (4-30Hz) or 1Hz-steps (31-91Hz) and converted to dB.

### Representational similarity analysis

Neural representational dissimilarity matrices (RDMs) were created from broadband responses (across time) or spectral responses (across frequencies). Neural RDMs were created in an analogous way to our previous studies on face and scene attractiveness [1,4], using the CoSMoMVPA toolbox for Matlab [14]. For the broadband responses, analysis was done separately for consecutive time-bins of 50ms, where all raw data falling into each time bin was used as a composite response pattern across sensors and time. For the spectral responses, analyses were done separately for each frequency step. The data of each participant was repeatedly split into two halves (each possible 50/50 split of the four runs). The first half of the data was used for dimensionality reduction of the neural response patterns [15]: The response patterns across channels was subjected to a PCA and the components explaining 99% of the variance were retained. The PCA solution was projected onto the second half of the data. The second half of the data was then used to compute neural dissimilarity: The response patterns were correlated (Spearman-correlations) across all possible pairs of videos, yielding a 75 × 75 neural RDM for each timepoint (in the analysis of broadband responses) or each frequency (in the analysis of spectral responses). These neural RDMs were then correlated (Spearman-correlations) with a set of model RDMs, separately for each participant; for these correlations, only the lower off-diagonal entries of the matrices were used.

I devised four model RDMs (Figure 1c). First, I created an RDM that reflected the videos’ pairwise dissimilarities in beauty ratings. For this RDM, the absolute difference between each participants’ own beauty ratings (averaged across the four repetitions) was computed for each pair of videos. Each participant’s neural data was thus modelled using their own beauty ratings. Second, I created an RDM that reflected the videos’ pairwise dissimilarities in category. For this RDM, scenes from the same category (e.g., two videos depicting animals) were coded as similar (0) and two videos from different categories were coded as dissimilar (1). Third, I created an RDM that reflected the videos’ pairwise dissimilarities in high-level features extracted by a GoogLeNet deep neural network [16] trained on scene categorization using the Places365 image set [17]. Previous studies showed that representations emerging in late layers of categorization-trained DNNs are predictive of high-level scene representations in the human brain, both in M/EEG and fMRI data [18–20], rendering them a good model for the extraction of the complex features diagnostic for visual scene attributes. For the deepnet RDM, activations were extracted from the last layer of the DNN and correlated for each pair of images; correlations were then subtracted from 1. Fourth, I created an RDM that reflected the videos’ pairwise dissimilarities in low-level visual motion energy measured by a wavelet-pyramid-based model [21]. Previous studies showed that this model predicts early visual responses to dynamic visual stimuli [21,22], rendering it suitable for modelling time-varying low-level features in the videos. For the motion energy RDM, model activations were extracted and correlated across each pair of videos; correlations were then subtracted from 1.

For the control analyses, I re-computed the correlations of the neural RDMs and the model RDM based on beauty ratings while partialing out (using partial Spearman-correlations) the model RDMs based on category, deepnet features, and motion energy.

For the broadband responses, all correlations between the neural RDMs and model RDMs were smoothed with a running average of 5 time points (i.e., 250ms), separately for each participant.

### Statistical testing

One-sided t-tests were used to compare correlations against zero. One-sided tests are chosen, because only correlations above zero are meaningful in RSA; negative correlations can only arise by chance or from misspecifications of representational models. P-values were FDR-corrected for multiple comparisons; only effects with p_corr_<0.05 are reported.

### Data sharing

Data are available on OSF (https://doi.org/10.17605/OSF.IO/5HYZN). Other materials are available upon request.

## Results

To quantify the neural correlates of aesthetic perception, I computed pairwise neural distances between all videos used in the experiment in a representational similarity analysis (RSA). First, I computed similarities between videos from correlations of evoked broadband responses across MEG sensors, in steps of 50ms (see Materials and Methods). I then modelled these neural distances with a predictor that captured how similar the videos were rated by the participants (Figure 1c); for details on the comparison between the predictor and the neural data, see Materials and Methods. This analysis revealed sustained representation of aesthetic quality across the whole video presentation, starting from 250-300ms (peak at 6,000-6,050ms) (Figure 1d).

To unequivocally attribute these effects to differences in aesthetic perception, I devised a set of three control predictors, which captured (i) whether the videos belonged to the same or to different visual categories, (ii) how similar the videos were in the high-level features extracted by the final layer of deepnet trained on scene categorization, and (iii) how similar the videos were in their spatiotemporal energy in low-level visual features (Figure 1c; see Materials and Methods). These control predictors also reliably correlated with neural representations (Figure 1d).

To assess whether information contained in the videos’ category, deepnet features and low-level motion energy could explain the neural correlates of aesthetic judgments, I repeated the analysis while partialing out the control predictors. In this analysis, aesthetic quality ratings still predicted neural responses in a reliable fashion from 500-550ms (peak at 6,700-6,750ms) (Figure 1e), suggesting that the sustained representation of beauty was not driven by properties captured by the control predictors.

To unveil the spectral correlates of aesthetic perception, I performed an RSA on trialwise powerspectra extracted from ongoing activity during the video presentation (see Materials and Methods). Pairwise similarities between videos were constructed from correlations between power values across all MEG sensors, separately for each frequency. I then again modelled these using the four predictors used above. Ratings of aesthetic quality were reliably associated with spectral features in the alpha (9-9.5Hz, peak at 9.5Hz) and beta (16.5Hz, 19-21Hz, 25.5Hz, peak at 20Hz) frequency bands (Figure 1g), also when partialing out the control predictors (alpha: single peak at 9.5Hz; beta: 16.5Hz, 19-20.5Hz, 25.5Hz, peak at 20Hz) (Figure 1f).

To assess whether the spectral signatures of aesthetic perception obtained previously were due to the frequencies contained in the visual response evoked by the video onset, we conducted an analysis in which we only used the data segments from 2-4s, 4-6s, and 6-8s. The analyses details were otherwise identical. This analysis revealed significant correspondences between the beauty ratings and neural representations in the alpha (8-10.5Hz, 12Hz, peak at 9.5Hz) and beta bands (16-17.5Hz, 19-20.5Hz, 24-24.5Hz, 25.5Hz). A similar result was obtained when partialing out the control predictors (alpha: single peak at 9.5Hz; beta: 16.5-17Hz, 19.5Hz, 20.5Hz, 24-24.5Hz, peak at 24.5Hz). This shows that the spectral alpha/beta signature obtained previously (Figure 1e/g) is not solely explained by frequencies contained in the initial evoked response.

Finally, a detailed analysis of the behavioral beauty ratings revealed significant differences in ratings across categories (one-way ANOVA: F[4,76]=4.35, p<0.001). To test whether these differences between categories account for the neural correlates of aesthetic ratings, I performed an analysis where I correlated the neural RDMs with the RDM based on beauty ratings while partialing out a predictor RDM that reflected the categories’ absolute differences in mean ratings. For the broadband responses, I still obtained a significant correlation that was highly similar to the previous results (Figure 1d/e), subtending large parts of the epoch from 500-550ms (peak at 6,000-6,050ms). For the spectral responses, I again found correspondences in the alpha (single peak at 9.5Hz) and beta (16.5Hz, 17.5Hz-18Hz, 19-21Hz, 25.5Hz, peak at 20Hz) bands. This shows that the effects cannot be accounted for by mean rating differences between categories.

Together, the current findings reveal a sustained neural representation of beauty for natural and dynamic visual inputs and establish a spectral neural signature of aesthetic perception in the alpha/beta frequency range.

## Discussion

The current results demonstrate that the aesthetic appeal of dynamic natural videos is not only encoded in sustained broadband responses, but also in spectral activations in the alpha and beta frequency range. Control analyses further suggest that the spectral coding of aesthetic appeal is independent of scene category, complex deepnet features, and low-level motion energy. These findings provide a spectral signature of aesthetic perception under conditions that are more ecologically valid than previously used static images, as they better mimic the dynamic visual experiences we encounter in real life.

Two previous studies have looked at spectral neural correlates of aesthetic perception with static images, suggesting that rhythmic neural codes are important for extracting aesthetic appeal for both moving and static inputs. First, consistently with our study, Kang and colleagues [2] reported an association between facial beauty and EEG alpha activity. Second, Strijbosch and colleagues [23] reported that gamma activity differentiated between aesthetically moving and non-moving art, while beta activity differentiated between moderately and highly moving art. My results are consistent with the low-frequency effects reported in this study. The absence of gamma-band differences here may be related to the longer presentation times or the everyday videos not spanning a sufficient range of beauty ratings. To test the latter assertion, future M/EEG studies need to present artworks and natural scenes within the same experiment, as in separate experiments participants independently adjust the range of their ratings to the range of aesthetic appeal encountered in each experimental setting.

Comparing the current results with studies using static stimuli further prompts the question whether neural rhythms are similarly associated with extracting aesthetic appeal for static and dynamic inputs. There are two reasons to expect that rhythmic neural correlates of beauty are more efficiently retrieved with moving stimuli: First, moving stimuli have been reported to be more engaging and to lead to stronger aesthetic experiences [7], which may enhance neural representations of beauty, too. Second, responses to static stimuli are largely driven by evoked visual responses, so that spectral analyses on such data may be driven by frequency components contained in the evoked response. Here, I show that aesthetic appeal is also encoded in ongoing rhythms when the strong initial evoked response is discarded; such evidence is harder to provide for static stimuli. Future studies need to include both static and dynamic stimuli to establish whether rhythmic brain activity predicts aesthetic perception similarly across these inputs.

A possible explanation for the beauty-related alpha/beta activations is their putative role in top-down processing [12]: Under conditions of sustained real-world experiences, it is likely that top-down modulations, caused by cognitive evaluations [24], play a major role in determining aesthetic appeal. Though largely speculative at this point, this assertion could be tested in future studies combining time-resolved M/EEG with spatially resolved fMRI.

Together, the current study provides an important starting point for further consideration of spectral signal components in visual neuroaesthetics.

## Acknowledgements

MEG data was acquired at the York Neuroimaging Centre. Thanks to Richard Aveyard for help with MEG setup and acquisition. D.K. is supported by the DFG (SFB/TRR135 – INST162/567-1) and “The Adaptive Mind”, funded by the Excellence Program of the Hessian Ministry of Higher Education, Science, Research and Art.

